# Extending rnaSPAdes functionality for hybrid transcriptome assembly

**DOI:** 10.1101/2020.01.24.918482

**Authors:** Andrey D. Prjibelski, Giuseppe D. Puglia, Dmitry Antipov, Elena Bushmanova, Daniela Giordano, Alla Mikheenko, Domenico Vitale, Alla Lapidus

## Abstract

**Background:** *De novo* RNA-Seq assembly is a powerful method for analysing transcriptomes when the reference genome is not available or poorly annotated. However, due to the short length of Illumina reads it is usually impossible to reconstruct complete sequences of complex genes and alternative isoforms. Recently emerged possibility to generate long RNA reads, such as PacBio and Oxford Nanopores, may dramatically improve the assembly quality, and thus the consecutive analysis. While reference-based tools for analysing long RNA reads were recently developed, there is no established pipeline for *de novo* assembly of such data.

**Results:** In this work we present a novel method that allows to perform high-quality *de novo* transcriptome assemblies by combining accuracy and reliability of short reads with exon structure information carried out from long error-prone reads. The algorithm is designed by incorporating existing hybridSPAdes approach into rnaSPAdes pipeline and adapting it for transcriptomic data.

**Conclusion:** To evaluate the benefit of using long RNA reads we selected several datasets containing both Illumina and Iso-seq or Oxford Nanopore Technologies (ONT) reads. Using an existing quality assessment software, we show that hybrid assemblies performed with rnaSPAdes contain more full-length genes and alternative isoforms comparing to the case when only short-read data is used.

**Availability and implementation:** rnaSPAdes is implemented in C++ and Python and is freely available for Linux and MacOS under GPLv2 license at cab.spbu.ru/software/rnaspades/ and github.com/ablab/spades.

## 1 Background

While a significant fraction of transcriptome studies are based on reference-assisted RNA-Seq analysis, they are limited to species with high-quality reference genome sequences and gene annotation. *De novo* assembly of RNA-Seq data allows to analyse transcripts sequences and gene expression without a reference genome, thus presenting a viable alternative to reference-based approaches. However, due to the short length of Illumina reads, recovery of complete transcript sequences originated from complex isoforms appears to be impossible without additional information, e.g. a reference genome. Recent biotechnological advances allowed to apply long-read technologies to transcriptome sequencing [1,2]. While long RNA reads seem to be extremely promising for transcriptomic studies, there is clearly a lack of software developed for their analysis.

In the past few years, several research projects involving long-read RNA sequencing were carried out [3–7]. All of them, however, were performed for species with relatively well-sequenced genomes. In these studies, researchers used such reference-based tools as Iso-seq pipeline [2], or designed in-house pipelines based on previously developed spliced aligners and tools for short-reads data analysis. However, no studies involving *de novo* assembly of long RNA reads were reported.

Most of the existing *de novo* transcriptome assemblers do not support long error-prone reads, since they were designed specifically for short Illumina reads. Among these tools only Trinity [8] supports hybrid assembly using corrected long reads. The only tool specially designed for hybrid transcriptome assembly is IDP-denovo [9], which is capable of improving third-party assemblies using long uncorrected reads. In addition, according to its manual, recently developed RNA-Bloom assembler [10] is capable of performing assembly solely from RNA ONT reads.

In this work we propose an extension for rnaSPAdes *de novo* transcriptome assembler [11]. Combining rnaSPAdes with previously developed hybridSPAdes approach [12] allows to exploit Iso-seq and ONT RNA reads as additional input and perform hybrid assembly. Since long-read technologies have a beneficial feature of detecting full-length (FL) mRNA sequences using terminal adapters in raw reads, a new version of rnaSPAdes can additionally take FL reads as an input, which further helps to determine complete isoform sequences.

To benchmark the assembly software, we selected several datasets containing both short and long reads. Although a variety of publicly available long-read RNA sequencing data is relatively small compared to conventional RNA-Seq, for this publication we selected three human datasets. The human transcriptome contains complex alternative isoforms, which allows to show the impact of using long reads for the assembly. Additionally, it eases the assembly quality evaluation, since the human genome is comparably well annotated and the ground truth is known. To assess generated assemblies we used rnaQUAST [13], which allows to evaluate their correctness and completeness using reference genome and gene database. Performed benchmarks show that incorporating long reads into the assembly pipeline allows to accurately assemble more complete genes and isoforms.

## 2 Methods

SPAdes assembly pipeline [14] consists of the four major steps: (i) de Bruijn graph construction from short reads, (ii) graph simplification, which removes erroneous edges from the graph and produces a so-called assembly graph, (ii) alignment of paired reads to the assembly graph and (ii) repeat resolution and scaffolding in the exSPAnder module [15, 16].

HybridSPAdes [12] additionally includes mapping long error-prone reads using BWA MEM algorithm [17] and exploiting these alignments during repeat resolution stage. Since hybridSPAdes is designed for genomic data, it heavily relies on unique (non-repetitive) edges in the assembly graph, which are selected using coverage and length criteria. An edge is considered to be unique if it has coverage close to the average coverage of the dataset and its length exceeds a certain threshold [12]. Indeed, such heuristics is not applicable for transcriptomics data, where the majority of edges are short and coverage is non-uniform.

In rnaSPAdes, graph simplification is modified specifically for RNA-Seq data and the repeat resolution step is substituted with an isoform reconstruction procedure [11]. However, the current version of rnaSPAdes is capable of using only short paired-end and single reads. To extend its functionality for hybrid transcriptome assembly, we combine it with procedures implemented in hybridSPAdes (see Fig. 1). While the read mapping step for transcriptomic data remains unmodified (with the exception of some alignment parameters), alterations were introduced to the isoform reconstruction procedure.

**Figure 1:**
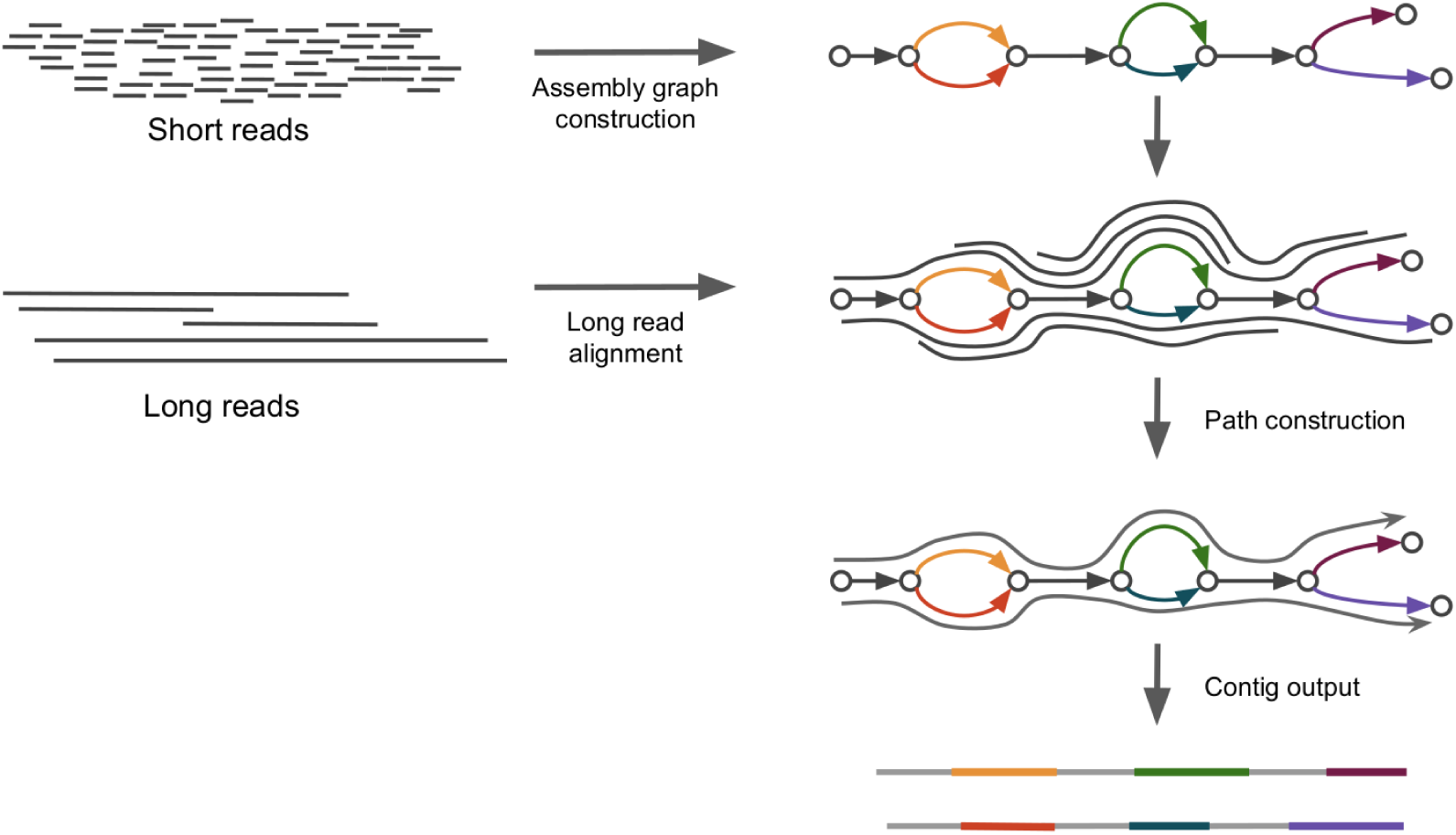
Hybrid rnaSPAdes pipeline. The assembly graph is constructed and simplified using only accurate short reads. Long error-prone reads are aligned to the graph. The resulting alignments are used in isoform reconstruction procedure. Grey edges represent common exons, colored ones correspond to alternative exons.

Similarly to genomic SPAdes, in rnaSPAdes isoform reconstruction is based on the concept of path extension implemented in the exSPAnder module. During path prolongation exSPAnder uses all available information simultaneously. In case of hybrid assembly, at every step exSPAnder tries to find correct extension edge using paired-end reads first, and then applies long-read path extension only if paired-end reads do not help (see [12, 15] for details).

Since alternatively spliced isoforms may form very similar paths, e.g. differing only by a single alternative exon, the key modification introduced to the path-extension procedure compared to the genomic pipeline is the possibility to select more than a single extension edge at each step. The same idea can be used for exploiting long-read alignments during the isoform reconstruction stage.

To extend a path *P* = (*p*_1_, *…, p*_*n*_) the algorithm considers all long-reads paths matching with *P*. A path *R* obtained from a long read alignment is defined as matching with *P* if there exists a suffix of *P* that is a prefix of *R*, or *P* is contained inside *R* (Fig. 2a). Formally, either (i) *R* = (*p*_*i*_, *…, p*_*n*_, *x*_1_, *…, x*_*k*_), *i* >= 1 or (ii) *R* = (*r*_1_, *…, r*_*l*_, *p*_1_, *…, p*_*n*_, *x*_1_, *…, x*_*k*_), where *r*_1_, *…, r*_*l*_ and *x*_1_, *…, x*_*k*_ are arbitrary edges in the graph. Further, from a set of all matching long-read paths the algorithm selects only those, for which the longest common subpath with *P* is (i) at least *L*_*min*_ long and (ii) contains at least *N*_*min*_ edges (default parameters are *L*_*min*_ = 200 *bp* and *N*_*min*_ = 2). The final set of matching long-read paths is denoted as **R_P_**. Then, among the set of all possible extension edges {*e*1, *…, e*_*m*_}, the algorithm selects all *e*_*i*_, such that at least one path from **R_P_** matches (*p*_1_, *…, p*_*n*_, *e*_*i*_) (Fig. 2b). Using only paths from **R_P_** instead of all matching long-read paths prevents from selecting all possible extensions for path *P*.

**Figure 2:**
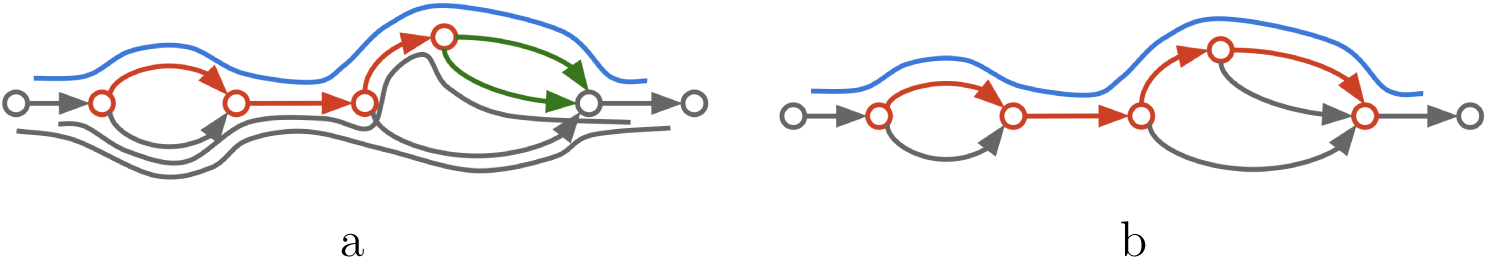
Example of path extension procedure using long reads. Red edges represent a current path being extended, green — possible extension edges, lines along the graph — long reads aligned to the graph. (a) Among all aligned reads the algorithm select only those that comply with the current path (blue line). (b) Complying paths are being used to extend the current path.

Paths in the graph are iteratively extended using paired-end and long reads until every edge is included in at least one path. Finally, to exploit reads capturing full-length transcripts, rnaSPAdes aligns them to the graph and produces FL paths, which are directly added to the set of resulting paths. Identical paths and exact subpaths are removed to avoid duplications, and the resulting set of paths is outputted in FASTA format.

## 3 Results

In this manuscript, we present quality reports for the assemblies of three datasets containing short and long-read sequencing data. Details on the used data are provided in Table 1. All datasets were quality-checked with FastQC [18] and trimmed with Trimmomatic [19] when adapters or significant quality drop were detected.

**Table 1:** Human transcriptome datasets selected for benchmarking.

| Dataset | Technology | # of reads | Strand-specific | Accession # |
| --- | --- | --- | --- | --- |
| Human | Illumina | 117 M | RF | SRP126849 |
| siNT_48 | Iso-seq | 5.6 M | — | SRP126849 |
| Human | Illumina | 84 M | No | SRX426377 |
| MCF7 | Iso-seq | 1.8 M | — | — |
|  | Iso-seq FL | 0.5 M | — | — |
| Human | Illumina | 54 M | No | SRR3103887 |
| GM12878 | Nanopores | 10 M | — | — |

We ran rnaSPAdes and Trinity with default parameters on Illumina data alone and on combined datasets. Since Trinity supports only corrected long reads, they were corrected using Illumina reads prior to the assembly with Racon [20]. Unfortunately, on hybrid data Trinity pipeline ran for over 4 weeks and did not produce the assembly. To perform hybrid assembly with IDP-denovo, we provided rnaSPAdes contigs and long raw reads as an input. However, either the resulting assembly remained unchanged, or IDP-denovo crashed. To compare hybrid transcriptome assembly against long-read-only assembly, we launched the only available tool RNA-Bloom on long reads from all 3 datasets (although it was designed only for ONT reads). Unfortunately, in all 3 cases RNA-Bloom managed to perform only read error correction step, but never completed the assembly itself. Thus, we present only results for Trinity and rnaSPAdes on short reads, and rnaSPAdes on hybrid data. Additionally, to evaluate the effect of external read error correction we ran rnaSPAdes on the same hybrid datasets, but using long reads corrected by Racon and RNA-Bloom. For a fair comparison, the same minimum contig length threshold was used for all assemblies (200 bp).

To evaluate the assembly quality we used rnaQUAST [13], which was designed specifically for assessing *de novo* transcriptome assemblies of organisms with high-quality reference genome and gene annotation. Among the large variety of metrics reported by rnaQUAST, we have selected only those that represent the most important characteristics of assembled sequences. In our opinion, one of the most significant statistics is the number of X%-assembled genes/isoforms, i.e. that have at least X% of bases captured by a single assembled contig. This metric shows the ability of *de novo* assembler to reconstruct complete transcript sequences, i.e. that can be used for further analysis. Presented reports also include database coverage (percentage of reference transcriptome nucleotides covered by all contigs), duplication ratio (the total number of aligned bases in all contigs divided by the total number of covered bases in reference isoforms, 1.0 in an ideal case), number of misassemblies (e.g. chimeric contigs) and average number of mismatch errors per contig. More details on various metrics and methods for assessing quality of *de novo* transcriptome assembly can be found in [13, 21, 22]. All datasets were assessed using Ensembl *H.sapiens* GRCh38.82 reference genome and gene annotation.

Tables 2, 3 and 4 demonstrate short quality reports for 3 human datasets. Since rnaSPAdes typically generates assemblies with higher quality than Trinity, rnaSPAdes is taken as a baseline in the following comparisons. As quality reports indicate, using raw long reads approach allows to reconstruct 8.4% more 95%-assembled genes on average comparing to short-read assembly. More importantly, the increase in 95%-assembled isoforms is larger (13.5% on average), which emphasizes that long reads are not only capable of reconstructing complex gene sequences, but are also useful for detecting alternatively spliced isoforms. One may also notice that the highest increase in 95%-assembled genes and isoforms (16.9% and 26.9% respectively) is achieved on Human MCF7 dataset, which contains FL reads. This fact suggests that FL reads can, unsurprisingly, be very beneficial for transcriptome assembly. Since long error-prone reads are used only to detect paths in the graph, per base accuracy decreases only by 0.32 mismatches per assembled transcript on average when long reads are added.

**Table 2:** Comparison between Trinity and rnaSPAdes on short-read data, and hybrid assembly performed by rnaSPAdes using raw long reads and long reads corrected with Racon and RNA-Bloom for Human MCF7 dataset. Since for this dataset FL reads are available, they were fed into rnaSPAdes using the appropriate option. Best value in each row is highlighted with bold.

| Assembly | Trinity | rnaSPAdes | rnaSPAdes<br>Hybrid | rnaSPAdes+<br>Racon | rnaSPAdes+<br>RNA-Bloom |
| --- | --- | --- | --- | --- | --- |
| Transcripts | 305K | 243K | 264K | 260K | 270K |
| Misassemblies | 2719 | **2170** | 3968 | 3001 | 4059 |
| Mismatches / contig | 1.85 | **1.50** | 2.31 | 1.82 | 2.26 |
| Database coverage | **0.26** | 0.23 | 0.25 | 0.24 | 0.25 |
| Duplication ratio | 2.45 | **1.45** | 2.96 | 2.28 | 2.94 |
| 50%-ass. genes | 13422 | 13546 | **13990** | 13728 | 13974 |
| 95%-ass. genes | 6091 | 6310 | **7379** | 7061 | 7370 |
| 50%-ass. isoforms | **24575** | 21048 | 23885 | 22427 | 23795 |
| 95%-ass. isoforms | 7856 | 7355 | **9333** | 8657 | 9273 |

**Table 3:** Comparison between Trinity and rnaSPAdes on short-read data, and hybrid assembly performed by rnaSPAdes using raw long reads and long reads corrected with Racon and RNA-Bloom for Human siNT_48 dataset. Best value in each row is highlighted with bold.

| Assembly | Trinity | rnaSPAdes | rnaSPAdes<br>Hybrid | rnaSPAdes+<br>Racon | rnaSPAdes+<br>RNA-Bloom |
| --- | --- | --- | --- | --- | --- |
| Transcripts | 463K | 426K | 444K | 436K | 443K |
| Misassemblies | 2625 | **1978** | 2657 | 2389 | 2661 |
| Mismatches / contig | 1.78 | **1.32** | 1.43 | 1.39 | 1.43 |
| Database coverage | **0.29** | 0.25 | 0.26 | 0.26 | 0.26 |
| Duplication ratio | 2.43 | **1.49** | 2.08 | 1.88 | 2.07 |
| 50%-ass. genes | **15681** | 15573 | 15570 | 15549 | 15573 |
| 95%-ass. genes | 7516 | 7831 | **8186** | 8117 | 8179 |
| 50%-ass. isoforms | **29150** | 24411 | 25660 | 24965 | 25669 |
| 95%-ass. isoforms | 9817 | 9300 | **10038** | 9886 | 10023 |

**Table 4:**
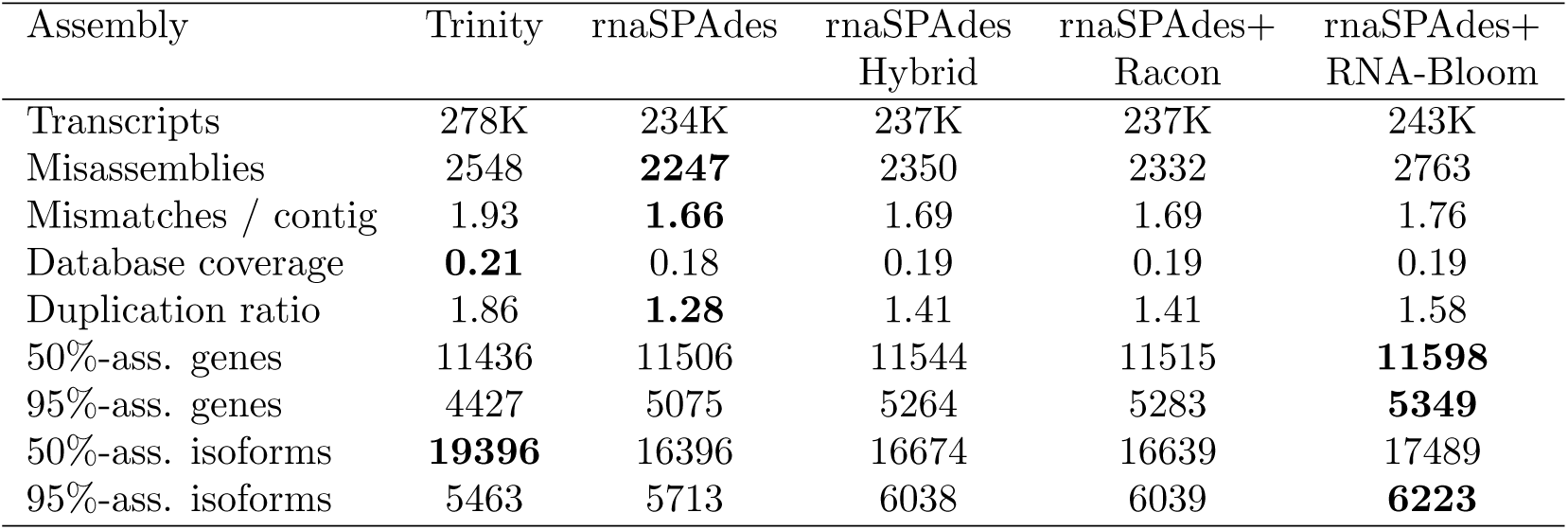
Comparison between Trinity and rnaSPAdes on short-read data, and hybrid assembly performed by rnaSPAdes using raw long reads and long reads corrected with Racon and RNA-Bloom for Human GM12878 dataset. Best value in each row is highlighted with bold.

Using pre-corrected Iso-seq reads, however, does not bring any substantial improvement comparing to raw data. Racon seems to remove a significant fraction of input data, which decreases the number of assembled genes and isoforms. Using RNA-Bloom’s corrected Iso-seq reads produces the assembly of almost the same quality. However, using ONT reads from Human GM12878 dataset corrected by RNA-Bloom allows to obtain slightly more 95%-assembled genes and isoforms comparing to the original data (1.6% and 3.1% respectively).

Exploiting long reads, however, also increases the number of misassemblies and duplication ratio. Manual analysis of misassembled contigs revealed that they are usually caused by the (i) presence of intronic and intergenic sequences, (ii) potentially unknown isoforms, (iii) merged neighboring genes and (iv) inaccurately assembled contigs that map to different loci, e.g. paralogous genes or intergenic space. However, since typical pipelines for analysis of transcriptomic data involve additional steps, such as transcript validation and annotation, duplicated and chimeric sequences have a less dramatic impact on the final results than, for example, in *de novo* genome assembly projects.

## Conclusion

In this work, we present a new rnaSPAdes workflow designed to improve *de novo* transcriptome assembly using long RNA reads. By using several human datasets containing both short and long reads, we show that the hybrid approach allows to restore more known genes, comparing to short-read only assemblers, Trinity and rnaSPAdes. We also demonstrate that long reads (especially FL-reads) can also be beneficial for discovery of alternatively spliced isoforms, which can be useful in various studies involving transcriptome analysis of previously unsequenced organisms.

## Availability of data and materials

Source code and manual of rnaSPAdes tool is available on cab.spbu.ru/software/rnaspades/.

Oxford Nanopore reads from Human GM12878 dataset are available at github.com/nanopore-wgs-consortium/NA12878/blob/master/RNA.md.

Iso-seq data for Human MCF7 dataset was released here datasets.pacb.com.s3.amazonaws.com/2015/IsoSeqHumanMCF7Transcriptome/list.html.

Other data is available at short read archive (ncbi.nlm.nih.gov/sra) with the following accession numbers

- Human GM12878: SRR3103887
- Human siNT _48: SRP126849 (siNT _48 sample)
- Human MCF7: SRX426377

## Acknowledgments

For uploading their data to public databases the authors would like to thank Nick Loman and other contributors of Oxford Nanopore RNA sequencing project, as well as the staff of following organizations: Stanford University, Netherlands Cancer Institute, Berlin Institute for Medical Systems Biology, and Pacific Biosciences.

